# Data-driven modelling of mutational hotspots and *in-silico* predictors in hypertrophic cardiomyopathy

**DOI:** 10.1101/826164

**Authors:** A.J. Waring, A.R. Harper, S. Salatino, C.M. Kramer, S Neubauer, K.L. Thomson, H. Watkins, M. Farrall

## Abstract

**Background:** Although rare-missense variants in Mendelian disease-genes have been noted to cluster in specific regions of proteins, it is not clear how to consider this information when evaluating the pathogenicity of a gene or variant. Here we introduce methods for gene-association and variant-interpretation that utilise this powerful signal.

**Methods:** We present a case-control rare-variant association test, *ClusterBurden*, that combines information on both variant-burden and variant-clustering. We then introduce a data-driven modelling framework to estimate mutational hotspots in genes with missense variant-clustering and integrate further *in-silico* predictors into the models.

**Results:** We show that *ClusterBurden* can increase statistical power to scan for putative disease-genes, driven by missense variants, in simulated data and a 34-gene panel dataset of 5,338 cases of hypertrophic cardiomyopathy. We demonstrate that data-driven models can allow quantitative application of the ACMG criteria PM1 and PP3, to resolve a wide range of pathogenicity potential amongst variants of uncertain significance. A web application (*Pathogenicity_by_Position*) is accessible for missense variant risk prediction of six sarcomeric genes and an R package is available for association testing using *ClusterBurden*.

**Conclusion:** The inclusion of missense residue position enhances the power of disease-gene association and improves rare-variant pathogenicity interpretation.

## INTRODUCTION

The clustering of pathogenic missense variants in specific regions or domains of proteins has been frequently reported [1–5]. A plausible mechanism underpinning this phenomenon is the presence of multiple loss or gain-of-function variants within functionally important domains [6]. Despite numerous examples of variant clustering, there have been few attempts to explicitly model variant residue position as a predictor of pathogenicity [7].

Pathogenic genes for Mendelian diseases were historically identified by linkage and candidate gene studies in multiple affected families [8]. Scanning of large exome patient cohorts offers an alternative strategy to identify novel pathogenic genes and variants. The aggregated burden of variants in affected cases compared to healthy controls has proved to be a useful test to confirm the pathogenicity of candidate genes [9], as well as identify novel putative pathogenic genes [10]. However, for genes with non-uniform variant pathogenicity, including positional information, alongside burden, may provide an increase in power to detect undiscovered low-penetrance genes.

The American College of Medical Genetics and Genomics (ACMG) have produced guidelines to interpret variant pathogenicity [11]. These guidelines integrate diverse data and classify variants into five categories from benign to pathogenic. However, due to limited information available for many variants, they fall into the category; ‘variant of uncertain significance’ (VUS). Although positional information is covered by criteria PM1, there is a lack of evidence for mutational hotspots, resulting in underapplication of this criteria. Furthermore, although much work has gone into the development of *in-silico* prediction scores, alternative scores can be conflicting, leading to discordance amongst testing laboratories [12] and uncertainty in their application. However, wherever large patient cohorts are attainable, mutational hotspots and the uncertainty surrounding *in-silico* predictors can be directly estimated from the data.

Hypertrophic cardiomyopathy (HCM), a relatively common autosomal dominant disease (1 in 500 prevalence), is a major cause of heart disease in people of all ages [13] and a cause of sudden cardiac death. In our cohort, eight sarcomeric genes collectively provide firm molecular diagnoses for ~27% of HCM patients, with a further ~13% of patients carry a VUS in the same genes. It has been suggested that disease and gene-specific approaches are needed to improve interpretation [14] and guidelines have been produced for specific genes and/or disease areas [15–18]. HCM is common enough to provide the large datasets needed for these gene-specific and data-driven approaches.

Here we propose new statistical approaches to explicitly include variant residue position in rare-missense variant association and interpretation; *ClusterBurden* for association testing and generalized additive models (*GAMs*) for hotspot estimation and application of *in-silico* prediction algorithms. We apply these methods to a large cohort of 5,338 HCM patients and up to 125,748 gnomAD [19] population controls. We demonstrate that utilising positional information increases power to detect disease-gene associations and elucidate the clustering signals present in 34 cardiomyopathy genes. We then use *GAMs* to model residue position and *in-silico* predictors for six core sarcomeric genes.

## METHODS

### Patient cohorts and simulated data

Next-generation sequence data for 34 cardiomyopathy genes (S1 Table) were available from two large HCM cohorts (S1A Methods); 2,757 probands referred to the Oxford Medical Genetics Laboratory (OMGL) for genetic testing and 2,636 probands recruited to the HCMR project [20]. High-coverage exonic sequences were captured by target enrichment and sequenced on the MiSeq platform (Illumina Inc.). Joint bioinformatic processing of both datasets followed the Genome Analysis ToolKit version 4 best practice guidelines (S1B Methods). OMGL variants were confirmed by Sanger sequencing and HCMR variants were manually checked by inspection of BAM files.

The gnomAD population reference database was used as a control group, which includes variant frequency data based on up to 125,748 individuals. For both cases and controls, only missense variants with a gnomAD population maximum allele frequency of less than 0.0001 [9–10] were included. This excludes potentially common ancestry-specific variants that are unlikely to be pathogenic for HCM.

To determine the theoretical performance of *ClusterBurden*, synthetic data were generated using a forward-time simulator (S2 Methods) designed to imitate rare-variants in genes with discrete exonic regions of increased pathogenic potential. Six different scenarios were considered, combining three clustering scenarios (uniform, a single pathogenic cluster and multiple pathogenic clusters) and two protein lengths (500 and 1,000 amino acid residues). For each scenario, 10,000 synthetic datasets were generated with 5,000 cases and 125,000 controls, variants were filtered at a simulated control MAF of <0.0001.

### Detecting missense variant burden and clustering – ClusterBurden

Missense variants causing amino acid substitutions at specific positions in the linear protein sequence were numbered from 1 to *N*, where *N* is the length of the protein. In all positional analyses, we consider this number as the position of a missense variant. As the background distribution of variants may be non-uniform, a cluster of variants observed in affected cases is insufficient in isolation to determine relevance to disease. Therefore to detect disease-relevant clustering, distributions were compared between affected cases and unaffected controls.

We propose *BIN-test* to evaluate these distributional differences. First, the protein’s linear sequence of amino-acid residues is split into *k* bins of equal length for both cases and controls. A chi-squared two-sample test is applied to the resultant *k* × 2 contingency table of binned variant counts. In simple terms, the test assumes that for each bin, the relative frequency of observed variants in cases and controls is the same and is more significant depending on how many bins deviate from this expectation and by how much. We applied a *k* ~ *n*^2/5^ heuristic [21] to select the optimal number of bins (*k*) dependent on *n*, the total number of observed variants. We compared the performance of the *BIN-test* with two other tests that compare distributions between two-samples; Anderson-Darling (AD) [22] and Kolmogorov-Smirnov (KS) [23]. Power and type 1 error were calculated using the (*r+1)/(n+1*) estimator where ***r*** represents the number of simulated datasets with p-values less than 0.05 and ***n*** is the number of simulations [24].

Current methods to discover novel Mendelian disease genes focus on the burden of rare-variants in an affected cohort relative to controls. We propose an approach, *ClusterBurden*, which tests the joint hypothesis of an excess of rare missense variants *and* differential clustering, in case-control data. This was accomplished by combining the p-values from a burden test (Fisher’s-exact test) with the *BIN-test*. As there are no known examples of a protective burden of rare exonic variants in cardiomyopathy, here we only consider an excess burden in the case group making it a one-sided test. Fisher’s method [25] was then used to calculate the joint significance of the combined p-values. An important assumption of this method is that the contributing p-values are independent; this was assessed in simulated data by Spearman’s rank correlation test [26]. The performance of *ClusterBurden* was compared, in simulated data, to two published position-informed association methods: DoEstRare [7] and CLUSTER [27], and three position-uninformed methods: C-alpha [28], SKAT [29], WST [30]. Then, both *BIN-test* and *ClusterBurden* were then applied to the 34 cardiomyopathy gene-panel in our HCM-gnomAD case-control cohort.

### Hotspot estimation and *in-silico* predictor modelling using GAMs

To test the hypothesis that a variant’s position can improve pathogenicity interpretation, we considered gene-specific models of variant clustering in cases and controls. By combining information on gene-level burden and variant positions, these data-driven models estimate the regional burden across the linear protein sequence to quantify mutational hotspots. The models were fitted in the GAM framework, [31] implemented in the R package “mgcv” [32]. The outcome variable was disease status, so each model was unsupervised with respect to previous classifications of pathogenicity. The training data included all rare-missense variants in cases and controls, including known pathogenic variants in the control set. Therefore, this approach implicitly models incomplete penetrance and benign background variation, leading to unbiased estimates of variant odds-ratios.

GAMs, as an extension of the linear modelling framework, are designed to deal with non-linear relationships of unknown complexity, between explanatory variables (e.g. residue position) and the response variable (e.g. case-control status). When a relationship is potentially non-linear, it is represented by a smooth curve instead of a straight line. These curves are inferred automatically using restricted maximum likelihood, which reduce over-fitting by penalising excessive ‘wiggliness’.

Using this framework, we defined the structure of the *hotspot-model*, which models; carrier status (gene-level burden) and residue position (clustering). To incorporate gene-level burden, non-carriers must also be modelled. However, as variant-level features such as residue position are meaningless for non-carriers, so a nested model structure is required, whereby residue position is included *only* as an interaction with carrier status. Under these circumstances, the smoothed residue position term is multiplied by zero for non-carriers, excluding this undefined data from the model. The structure of the *hotspot-model* is as follows:

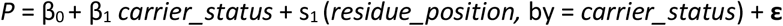

where **P** is the probability of being a case, **β_0_** is the model intercept, **β_1_** is a linear coefficient for *carrier_status*, **s_1_** is a smoothed (i.e. non-linear) function for *residue_position*, **by** is used to generate factor-smooth interactions and **ε** is a binomial error term.

The feasibility of this approach is dependent on the number of observations, thereby limiting its application in our data to the six core sarcomeric genes; *MYBPC3*, *MYH7*, *MYL2*, *MYL3*, *TNNI3* and *TNNT2*, each carrying at least 20 rare-missense variants. A *hotspot-model* was produced for each gene and raw model predictions for each residue position, in the form of logistic probabilities and standard errors, were transformed to ORs and 95% confidence intervals (CIs). There is currently no universal guidance on how to quantitatively apply ACMG criteria PM1. However, using the probability that a variant is a case variant as a proxy of pathogenicity, we can use predicted probabilities to attribute levels of evidence. Here we stratify variants based on the probability thresholds 0.9, 0.95 and 0.99 to represent supporting, moderate and strong evidence of pathogenicity. These correspond to ORs of approximately 10, 20 and 100.

As GAMs are additive in structure, it is straight-forward to include further predictors in the model. Here we experimented with the inclusion of variant prediction scores extracted from the dbNSFP4.0 database [33]. These *in-silico* prediction algorithms are covered by the ACMG criteria PP3, however, like criteria PM1, it is challenging to apply this criteria quantitatively. It is unclear which threshold determines a pathogenic variant with a given probability and whether these thresholds are consistent across genes. Both of these problems can be solved using gene-specific models as the relationship between the *in-silico* predictor and disease-status is automatically inferred. Furthermore, we can quantify uncertainty on the usage of these scores for each gene.

To avoid overfitting, we implemented a strict two-stage feature selection procedure, based on 24 *in-silico* scores from dbNSFP. In stage 1, only features with a marginal p-value < 0.002 (0.05/24 for Bonferroni correction) were selected. In stage 2, backwards elimination was implemented, whereby features are removed from the model one at a time and then remaining features are assessed again in a new model, until all features are significant (Bonferroni corrected for the number of features selected in stage 1). The resulting models, which are henceforth termed *in-silico-models*, assimilate evidence of gene-burden, variant clustering and pathogenicity prediction scores.

Model performance for these GAMs are best judged by the estimates of uncertainty accompanying predictions. However, to determine the relative ability of these models to predict pathogenicity, they were compared with models based on single *in-silico* predictors and expert variant classifications. Relative performance was assessed using the area under the curve (AUC) from the receiver operator characteristic (ROC) curves with ten cross-fold validations performed by dividing the data into training and test sets in the ratio of 80:20%.

## RESULTS

### Testing the hypothesis of clustered missense variants

Under the null hypothesis of no excess burden or clustering, the false-positive rate of the *BIN-test* and AD test were adequately controlled in simulated data, whereas the KS test was overly conservative (S3 Table). The *BIN-test* had superior power than AD or KS under all clustering scenarios and protein lengths with on average 1.8-fold more power to detect clustering. As power covaries with the number of observed variants, power was higher for longer proteins as well as proteins with more pathogenic residues.

Correlations between p-values generated by the *BIN-test* and Fisher’s exact test were compared for simulated data under 1) a null model of no association or 2) a disease model of over-burdened and clustered variants. For the disease model, there was a positive correlation (Spearman’s rank correlation rho=0.40) between p-values, as anticipated as the power of these tests covaries with the number of observed variants. However, under the null model, the p-values were completely uncorrelated (i.e. rho = 0.00), satisfying the independence assumption of Fisher’s method.

The false-positive rate for *ClusterBurden*, DoEstRare, CLUSTER and C-alpha were all well controlled in the simulated datasets (S3 Table). On the contrary, SKAT and WST showed markedly inflated false-positive rates under the null and were not examined further. *ClusterBurden* was the most powerful method when clustering was present with an average of 72% power, 3% higher than the second-best test DoEstRare. The best method under the uniform model (i.e. burden-only) was CLUSTER, which had ~5% more power than *ClusterBurden*. Amongst the position-informed tests, *ClusterBurden* was the most rapid to compute taking less than a second per gene whereas DoEstRare and CLUSTER took over 20 or 4 minutes respectively.

We then examined 34 cardiomyopathy genes for rare-missense variant associations with Fisher’s-exact test (burden), *BIN-test* (cluster) and *ClusterBurden* (combined cluster and burden) in our cohorts of HCM cases and gnomAD controls (Figure 1). Significance thresholds were conservatively Bonferroni adjusted to allow for 34 genes × 3 methods (i.e. p-values adjusted for 102 tests to p < 0.00049). Significant burden signals were then detected in 11 genes with Fisher’s-exact test; *MYH7* (p < 5.44 × 10^-252^), *MYBPC3* (p < 1.74 × 10^-229^), *TNNI3* (p < 1.46 × 10^-50^), *TNNT2* (p < 1.11 × 10^-24^), *TPM1* (p < 6.56 × 10^-21^), *ACTC1* (9.61 × 10^-14^), *GLA* (1.61 × 10^-10^), *FLH1* (1.02 × 10^-9^), *MYL2* (1.87 × 10^-9^), *CSRP3* (3.56 × 10^-8^) and *MYL3* (6.53 × 10^-6^). The *BIN-test* detected significant clustering for 6 core sarcomeric genes; *MYH7* (p < 1.36 × 10^-73^), *MYBPC3* (p < 1.55 × 10^-78^), *TNNI3* (p < 3.34 × 10^-13^), *MYL2* (p < 5.83 × 10^-10^), *TNNT2* (p < 1.69 × 10^-7^) and *MYL3* (p < 1.7 × 10^-4^). Two additional sarcomeric genes showed nominal evidence of clustering; *ACTC1* (p<0.0412) and *TPM1* (p< 0.0494). *ClusterBurden* confirmed the association for 11 genes that showed burden signals and calculated substantially lower p-values for the six core-sarcomeric genes with significant clustering, consistent with enhanced power for this approach in true disease-causing genes.

**Figure 1:**
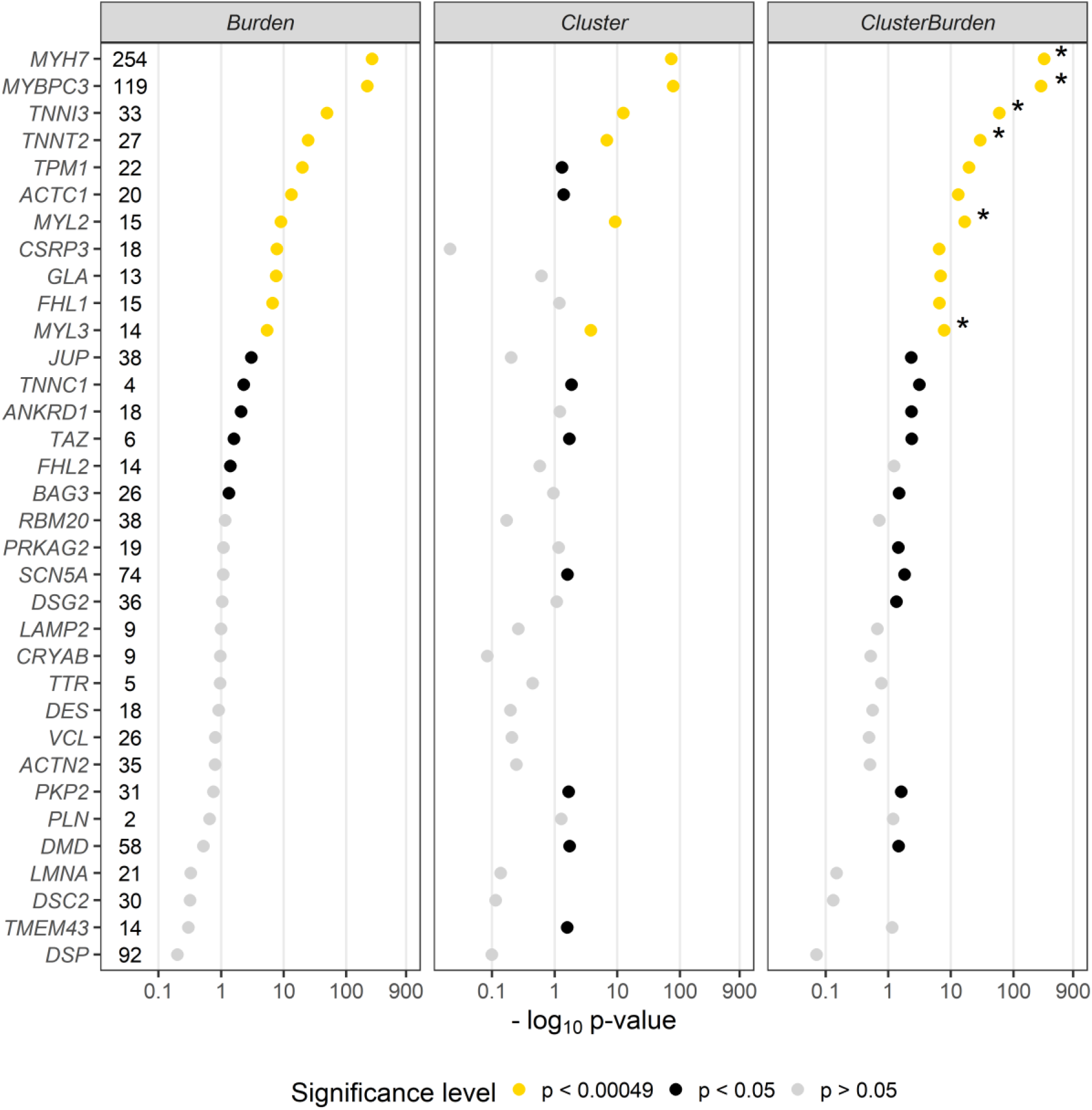
Association analysis using Fisher’s-exact test (Burden), *BIN*-test (Cluster) and *ClusterBurden* of 34 genes commonly present on cardiomyopathy gene-panels. Our case-control dataset contains 5,338 hypertrophic cardiomyopathy cases and 125,748 gnomAD controls. For all tests, only missense variants with a *popmax* MAF less than 0.01% were considered. P-values are presented on a −log_10_ scale. The number of observed case variants in each gene is displayed next to the gene symbol. P-values displayed in yellow are significant after Bonferroni correction for 34 genes × 3 tests (p < 0.00049), p-values in black are nominally significant (p < 0.05) and p-values in grey are insignificant (p > 0.05). Asterix’s denote genes where the *ClusterBurden* p-value is lower than the Burden p-value. Two vertical dotted lines at 0.05 and 0.00049 indicate the nominal and Bonferroni significance thresholds.

### Hotspot and in-silico models

Figure 2 summarises the *GAM-*predictions for six sarcomeric genes in the *hotspot-models*. Visualising the predicted odds-ratios (OR) for each residue illuminates the local burden of rare-missense variants across each protein, identifying “mutational-hotspots” and highlighting areas of potential functional importance in HCM pathogenesis. Confidence in these predictions is tight for *MYH7* and *MYBP3*, conversely the genes with fewer observed variants have much broader confidence intervals. ORs from all models correlate strongly with expert manually assigned classifications, though there is substantial overlap between classes. Variants with a VUS classification show the highest spread in predictions.

**Figure 2:**
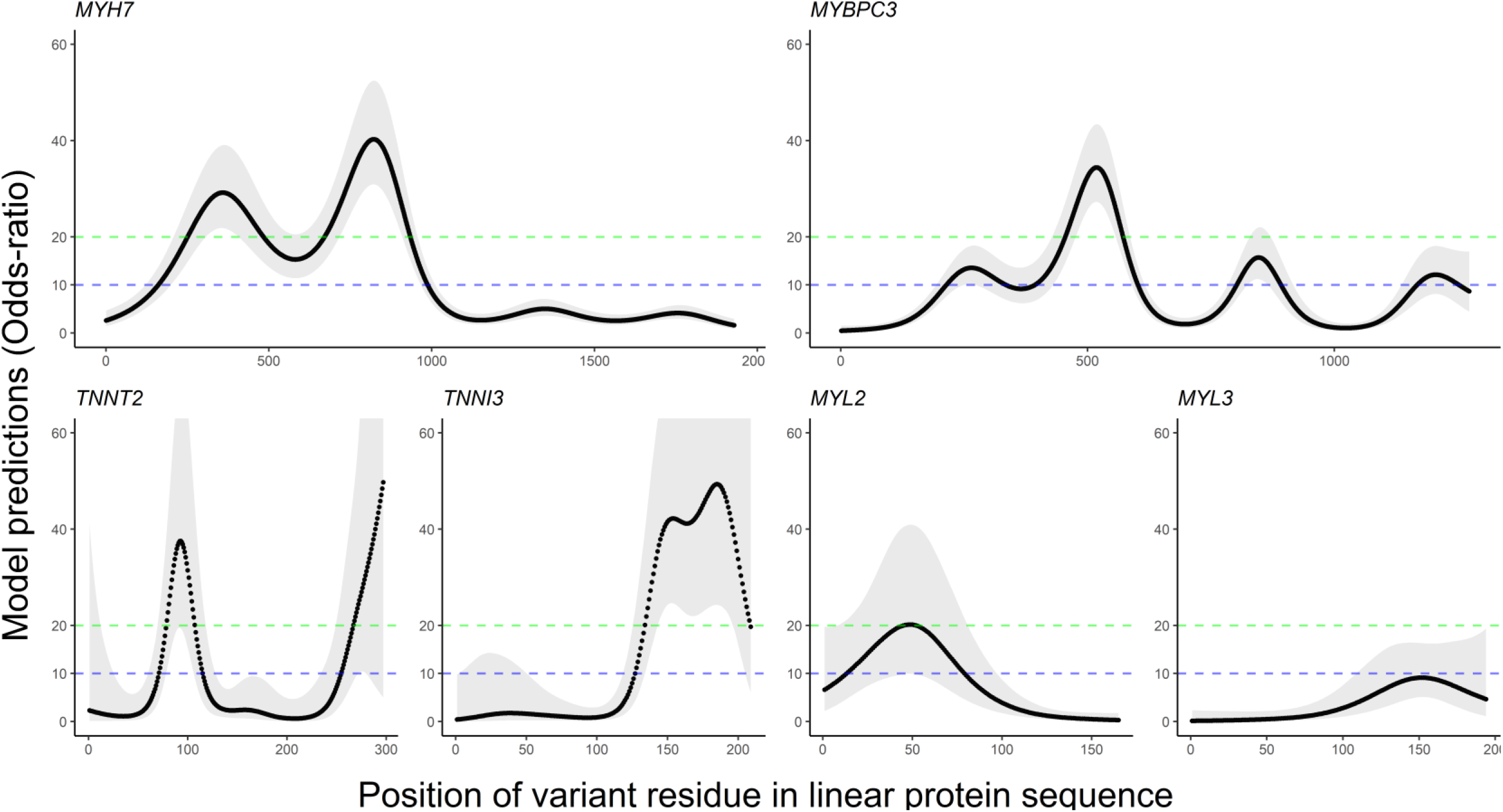
Odds-ratio (OR) predictions and 95% confidence intervals for *hotspot-models*. Mutational hotspots were estimated for six firmly established HCM disease-genes. The 95% confidence intervals for model predictions are displayed as light grey shading. Regions above the blue (OR=10) and green (OR=20) dashed lines ascribe supporting and moderate evidence respectively, for criteria PM1.

Figure 3 displays *in-silico-model* predictions for individual variants in the same six genes. Due to strict feature selection, the number of predictors included in each model depends on the power to detect associations between features and disease status. This resulted in fewer features for genes with fewer observed variants, *MYL3* had no additional significant features. As residue position is included as a predictor in each model, predictions generally follow the *hotspot-model*, however, due to additional *in-silico* predictors, ORs tend to vary from this pattern, stratifying risk for variants at the same position.

**Figure 3:**
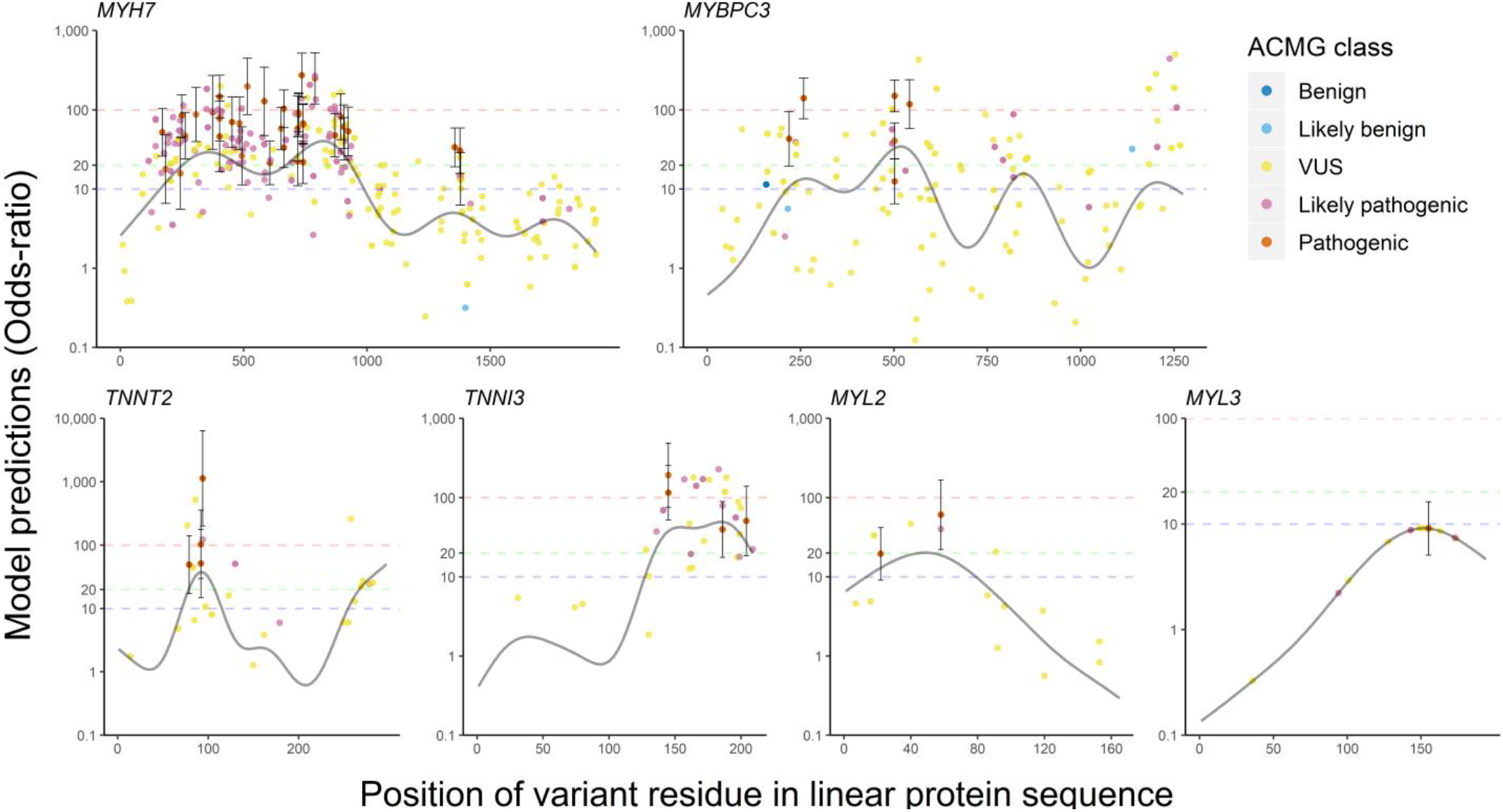
Odds-ratio (OR) predictions and 95% confidence intervals for *in-silico-models*. Each point denotes a rare-variant in the HCM dataset and is coloured to indicate its expert classification. ORs on the y-axis are displayed on a log_10_ scale and were derived from the *in-silico-models*, including; gene-burden, residue position and gene-specific significant secondary features from dbNSFP. The solid black curvy lines represent the predictions for each residue in the protein for a gene-burden and position model (*hotspot-model*). Dashed red lines indicate an OR of 1, dashed blue lines indicates the OR for the uniform burden model.

As with the *hotspot-models* there was strong correspondence between predictions and expert classifications (mean rho 0.41 across six models). In *MYH7*, mean predicted ORs for pathogenic, likely pathogenic and VUS variants observed in cases, were 74, 50 and 20 respectively. Again, the VUS class had highest heterogeneity, with predicted ORs ranging from 0.25 to 197 (*MYH7*). For half of these VUS’s, limited information is available, as they are observed in a single case and absent in controls (singleton variants). The empirical ORs for these singletons, based only on case and control frequencies and adding 0.5 to zero-count cells (Haldane continuity correction [34]), had very wide 95% Cis: 44.9 [1.5, 1338.3]. However, *predicted* ORs for such variants can have greater precision with different point estimates depending on the precise amino-acid substitution. In MYH7, five singleton VUS’s had predicted ORs greater than 100 and three had ORs less than 1.

The mean and standard deviation of AUC for ten cross-fold validations summarise overall model performance (Figure 4). The *in-silico-model* had a much higher mean AUC than any individual *in-silico* predictor in isolation. With the exception of *MYBPC3*, the *hotspot-model* out-performed any standalone score from dbNSFP. This suggests that residue position is more important in determining pathogenicity than the *in-silico* predictors in dbNSFP for these sarcomeric proteins. The AUC standard deviations for *MYH7* and *MYBPC3* were the smallest, suggesting they have the highest capacity to generalize to new variants.

**Figure 4:**
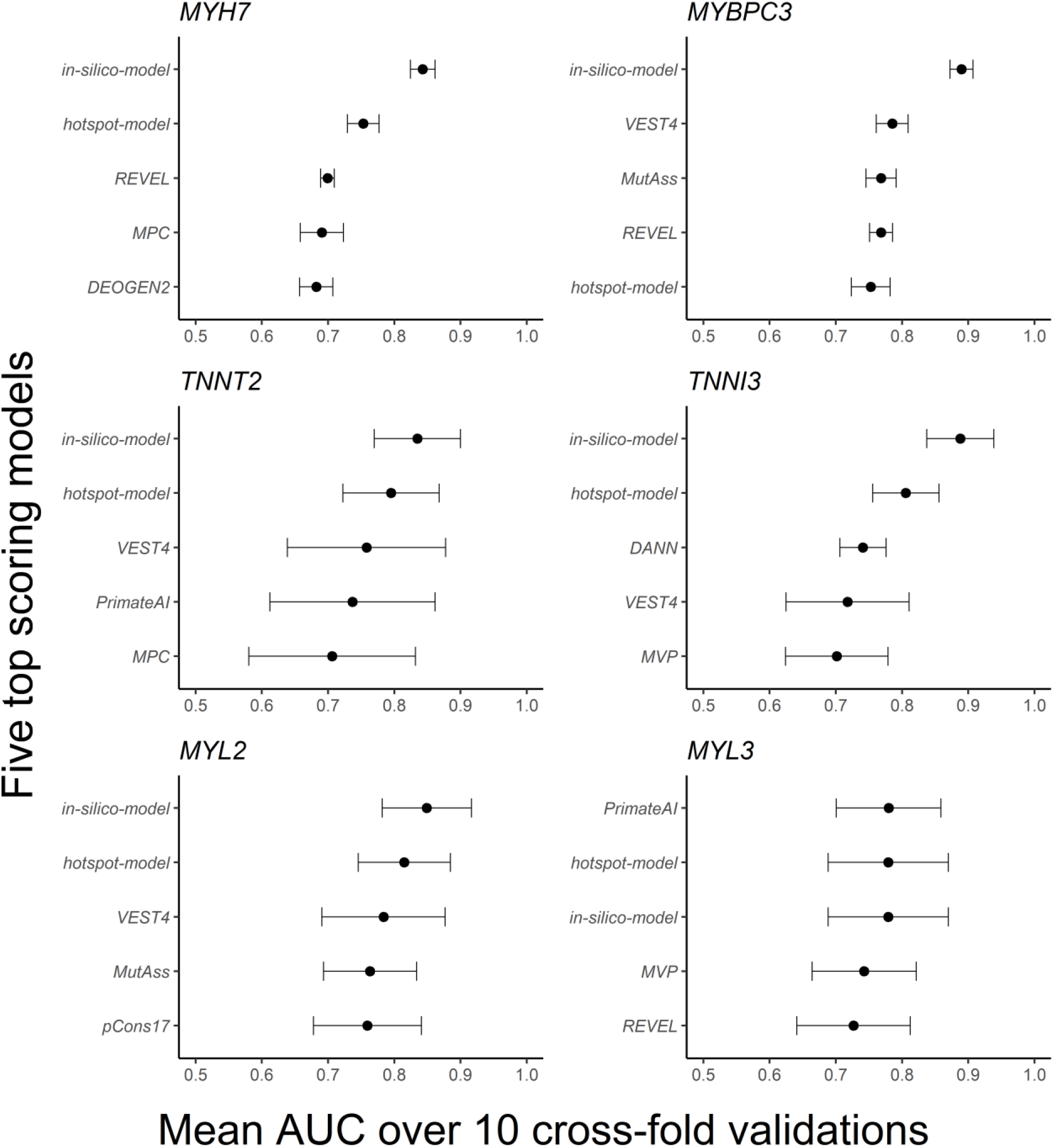
Means and standard deviations of area under the curve (AUC) across 10 cross-fold validations. For each gene, the *in-silico-model* and *hotspot-model* are compared to each individual *in-silico* predictor from dbNSFP. Each model is trained on the same HCM-gnomAD case-control missense variants all filtered at a gnomAD population maximum frequency of 0.01%. Only the five highest mean AUC scoring models are displayed.

The GAMs were then used to attribute evidence of pathogenicity based on the ACMG criteria PM1 and PP3. Using the ACMG OR thresholds described in the methods, Table 1 shows the proportion of variants in our cohort with evidence of pathogenicity predicted by the *hotspot* and *in-silico-models*. For the *hotspot-model*, the PM1 criteria was satisfied, with supporting (OR > 10) or moderate (OR > 20) evidence, for some variants in all genes except *MYL3*. Strong evidence of pathogenicity (OR > 100) was not predicted by the *hotspot-model* for any variant. Conversely, the *in-silico-model*, which combined criteria PM1 and PP3, provided strong evidence of pathogenicity for many variants, including VUS’s in *MYH7, MYBPC3, TNNT2* and *TNNI3*.

**Table 1:** Proportion of variants with evidence of pathogenicity in *hotspot* and *in-silico* models. Each observed variant across six genes in our HCM cases was given supporting (OR > 10; Sup.), moderate (OR > 20; Mod.) or strong (OR > 100; Str.) evidence of pathogenicity based on model predictions from the *hotspot* and *in-silico* models. The proportion of variants with supporting, moderate or strong evidence are stratified by expert classifications made by Oxford Medical Genetics Laboratory.

|  | **VUS** |  |  |  | **Likely pathogenic** |  |  |  | **Pathogenic** |  |  |  |
| --- | --- | --- | --- | --- | --- | --- | --- | --- | --- | --- | --- | --- |
|  | *N* | *Sup.* | *Mod.* | *Str.* | *N* | *Sup.* | *Mod.* | *Str.* | *N* | *Sup.* | *Mod.* | *Str.* |
| ***Hotspot-models*** |  |  |  |  |  |  |  |  |  |  |  |  |
| *MYH7* | **123** | 10% | 28% | 0% | **93** | 33% | 55% | 0% | **36** | 31% | 61% | 0% |
| *MYBPC3* | **97** | 35% | 12% | 0% | **13** | 38% | 23% | 0% | **6** | 33% | 67% | 0% |
| *TNNT2* | **19** | 21% | 47% | 0% | **4** | 0% | 50% | 0% | **4** | 0% | 100% | 0% |
| *TNNI3* | **18** | 17% | 67% | 0% | **11** | 9% | 91% | 0% | **4** | 0% | 100% | 0% |
| *MYL2* | **12** | 25% | 0% | 0% | **1** | 100% | 0% | 0% | **2** | 100% | 0% | 0% |
| *MYL3* | **10** | 0% | 0% | 0% | **3** | 0% | 0% | 0% | **1** | 0% | 0% | 0% |
| ***In-silico models*** |  |  |  |  |  |  |  |  |  |  |  |  |
| *MYH7* | **123** | 18% | 26% | 4% | **93** | 16% | 58% | 14% | **36** | 8% | 75% | 17% |
| *MYBPC3* | **97** | 16% | 25% | 7% | **13** | 15% | 54% | 15% | **6** | 17% | 33% | 50% |
| *TNNT2* | **19** | 16% | 26% | 16% | **4** | 0% | 50% | 25% | **4** | 0% | 50% | 50% |
| *TNNI3* | **18** | 22% | 33% | 22% | **11** | 18% | 45% | 36% | **4** | 0% | 50% | 50% |
| *MYL2* | **12** | 0% | 25% | 0% | **1** | 0% | 100% | 0% | **2** | 50% | 50% | 0% |
| *MYL3* | **10** | 0% | 0% | 0% | **3** | 0% | 0% | 0% | **1** | 0% | 0% | 0% |

A web application, *pathogenicity_by_postion*, is available to facilitate the exploration of the *hotspot* and *in-silico* models (R Shiny: https://adamwaring.shinyapps.io/Pathogenicitybyposition). Users can explore alternative models and submit their own missense variants to retrieve predicted ORs and support intervals. A further R package is available for cluster-detection and association testing using *BIN-test* and *ClusterBurden* (https://github.com/adamwaring/ClusterBurden).

## DISCUSSION

We present here new statistical approaches to incorporate residue position in the analysis of rare-missense variants in Mendelian disease genes. Our association tests were well-calibrated in simulated data, the *BIN-test* detected significant clustering in almost all firmly-established HCM genes, and *ClusterBurden* gave superior power over a simple burden test. Data-driven models were applied to six core sarcomeric genes to estimate mutational hotspots and provides a flexible method for quantitative application of ACMG criteria PM1 and PP3. Our results demonstrate that residue position can be a powerful predictor of both gene and variant pathogenicity and can quantify the statistical uncertainty surrounding the application of *in-silico* algorithms.

*BIN-test* is a powerful method to test the hypothesis of variant clustering in known or putative disease genes. *ClusterBurden* is a gene association test with superior power over a standalone burden test in situations where pathogenic variants cluster in specific protein regions. Both tests keep false-positives below 5% and are rapid to compute, making them scalable for whole-exome scanning of very-large datasets like the UK Biobank. Although *ClusterBurden* has slightly reduced power whenever clustering is absent, we observed clustering for most well-established HCM genes where missense variants cause disease. Therefore, this method has the potential be more powerful to detect undiscovered low-penetrance genes than a burden-only test.

The most significant position signal was observed in the beta myosin heavy chain protein (*MYH7:* ENST00000355349), a finding that has been long recognised [35]. Most of the signal was observed in the myosin-head motor domain between residues 100-900, in two clusters peaking at residues 370 and 830 (Figure 2). The relatively low variant density in cases and high density in controls in the carboxy-terminus of this protein might lead an observer to hypothesise a regional protective effect on HCM risk (S13 Figure). In sharp contrast, the GAM model predicts a modestly excessive burden (OR ~3) across this entire region discounting the likelihood of a localised protective effect (Figure 2).

A strong position signal, driven by four potential clusters, was observed in the *MYBPC3* gene (ENST00000545968), which encodes cardiac myosin-binding protein C C10 (Figure 2). Clusters peaked at residues 260, 518, 864 and 1274, which respectively fall in domains C1, C3, C7 and C10. Multiple functional roles are suspected for the region containing the C1 domain, including binding to myosin S2 and actin. The C10 domain is also a possible titin binding site [36]. To explore whether the signal was overly driven by high-frequency founder mutations, seven variants with allele counts above 10; c.2429G>A, c.2308G>A, c.1624G>C, c.1504C>T, c.1484G>A, c.772G>A and c.655G>C (ENST00000545968), were masked in a sensitivity analysis. In their absence, there is still strong evidence of a position signal (p < 3 × 10^-9^) and the remaining peak densities overlap with the locations of the (masked) founder mutations (S14 Figure).

Eighty-nine percent of 27 case variants in the *TNNT2* gene (ENST00000509001), which encodes cardiac troponin T, map to clusters between residues 67-179 and residues 250-282 (Figure 2). The first peak at residue 90 overlies a previously reported region at residues 79-179 that binds tropomyosin [37]. Mutations between residues 92-110 have been previously noted to impair tropomyosin dependent functions in *TNNT2* [38] and six variants map to this region. In *TNNI3* (ENST00000344887), which encodes cardiac troponin I, 91% of 34 case variants mapped to a cluster spanning residues 128-209. This accords with previous studies documenting disease-causing variant clustering in the carboxy-terminus of this sarcomeric protein [39]. In *MYL2* (ENST00000228841), which encodes myosin regulatory light chain, 50% of 30 case variants cluster between residues 25 and 100, whereas control variants tended to cluster towards the C-terminus (Figure 2). In *MYL3* (ENST00000395869), which encodes myosin essential light chain, 79% of 14 case variants cluster between residues 125 and 175 (Figure 2) whereas control variants were more uniformly distributed.

*GAMs* were used to model variant pathogenicity based on mutational hotspots (*hotspot-model)* and *in-silico* predictors (*in-silico-model*). GAMs have attractive statistical properties, not necessarily shared by other machine-learning approaches, in that they can produce familiar interpretable results via variant-specific ORs and accompanying 95% confidence intervals. Unlike empirical ORs, based solely on observed frequencies for variants, GAM ORs draw upon a much larger pool of information, including features of other variants in the training set. This permits the estimation of variant-specific ORs whenever the empirical frequencies are uninformative. Furthermore, as the GAM response variable is case status, the models are unbiased by previous classifications, and account for both incomplete penetrance and benign background rare-variation.

Reassuringly, model predictions were positively correlated with expert manually-curated classifications. Using a probabilistic approach, we attributed different levels of evidence for the criteria PM1 and PP3. Currently for HCM, criteria PM1 is applied as moderate evidence to *MYH7* for variants that fall in the residue 181-937 motor-domain [15]. The *hotspot-model* extends this criteria to five more sarcomeric genes and stratifies evidence as either supporting or moderate. When *in-silico* predictors were included in the model, evidence was stratified as supporting, benign or occasionally strong. However, this relies on effectively collapsing two ACMG criteria into one, a relevant modification of the current additive guidelines [11].

## CONCLUSIONS

Here we have demonstrated proof-of-concept data-driven modelling as an effective tool for variant interpretation in HCM. Collaboration for more cases could substantially improve model performance and reduce uncertainty in estimates, especially for genes with few observations such as *MYL2* and *MYL3*, but could also allow informative modelling of further HCM genes and other Mendelian diseases. Furthermore, the modelling approach developed could be modified into a Bayesian framework allowing inclusion of previously reported pathogenic and benign mutations, available from public resources such as clinVAR, as model priors, reducing uncertainty further. Our association methods, have proved theoretically powerful and could be used to scan for novel cardiomyopathy genes when large datasets such as 500K UK Biobank exomes become available.

## LIST OF ABBREVIATIONS

ACMG: American College of Medical Genetics
VUS: Variant of Uncertain Significance
HCM: Hypertrophic Cardiomyopathy
GAM: Generalized Additive Models
OMGL: Oxford Medical Genetics Laboratory
HCMR: Hypertrophic Cardiomyopathy Registry
BAM: Binary alignment map
AD: Anderson-Darling
KS: Kolmogorov-Smirnov
AUC: Area under the Curve
ROC: Receiver Operating characteristic
OR: Odds-ratio

## DECLARATIONS

### Ethics

The research protocol was approved by the South Central - Oxford A Research Ethics Committee (REC reference: 14/SC/0190); written informed consent was obtained from all participants.

### Availability of data and materials

Due to the confidential nature of some of the research materials supporting this publication not all of the data can be made accessible to other researchers. Please contact the corresponding author for more information.

### Author contributions

AW and MF led conception and design of work

AW contributed to data curation, conducted all analyses, developed statistical methods and associated software and wrote each draft of manuscript

MF supervised all analyses and interpretation of results

AH led the curation and quality control of the hypertrophic cardiomyopathy datasets

SS contributed to quality control of the data

CK and NS supported access to the HCMR dataset

KT and HW advised and guided the clinical aspects of the work

MF and KT reviewed and editing each draft of the manuscript

## Acknowledgements

We would like to acknowledge Anuj Goel for his bioinformatics support in data curation and Michael Bowman for his support in accessing data from the clinical genetics laboratory.

Computation used the Oxford Biomedical Research Computing (BMRC) facility, a joint development between the Wellcome Centre for Human Genetics and the Big Data Institute supported by Health Data Research UK and the NIHR Oxford Biomedical Research Centre. Financial support was provided by the Wellcome Trust Core Award Grant Number 203141/Z/16/Z. The views expressed are those of the author(s) and not necessarily those of the NHS, the NIHR or the Department of Health.

