## Supplementary Information for "Data-driven modelling of mutational hotspots and *in-silico* predictors in hypertrophic cardiomyopathy"

*S1 Table: The ensemble transcript ids for all genes considered in our analyses. For all genes, canonical ensembl transcripts were used.*

| *Gene* | *Transcript* |
|---|---|
| ACTC1 | ENST00000290378 |
| ACTN2 | ENST00000366578 |
| ANKRD1 | ENST00000371697 |
| BAG3 | ENST00000369085 |
| CRYAB | ENST00000533475 |
| CSRP3 | ENST00000533783 |
| DES | ENST00000373960 |
| DMD | ENST00000357033 |
| DSC2 | ENST00000280904 |
| DSG2 | ENST00000261590 |
| DSP | ENST00000379802 |
| FHL1 | ENST00000394155 |
| FHL2 | ENST00000358129 |
| GLA | ENST00000218516 |
| JUP | ENST00000393931 |
| LAMP2 | ENST00000434600 |
| LMNA | ENST00000368300 |
| MYBPC3 | ENST00000545968 |
| MYH7 | ENST00000355349 |
| MYL2 | ENST00000228841 |
| MYL3 | ENST00000395869 |
| PKP2 | ENST00000070846 |
| PLN | ENST00000357525 |
| PRKAG2 | ENST00000287878 |
| RBM20 | ENST00000369519 |
| SCN5A | ENST00000413689 |
| TAZ | ENST00000299328 |
| TMEM43 | ENST00000306077 |
| TNNC1 | ENST00000232975 |
| TNNI3 | ENST00000344887 |
| TNNT2 | ENST00000509001 |
| TPM1 | ENST00000358278 |
| TTR | ENST00000237014 |
| VCL | ENST00000211998 |

*S2 Table: Implementation details for all statistical tests used in the comparison of type 1 and 2 error.*

| *Test name* | *Implementation (all implemented in R)* | *Notes on implementation* | *Original paper* |
|---|---|---|---|
| Fisher's exact test | base::fisher.test | One-sided p-value (*alternative="greater")* | (Fisher 1925) |
| BIN-test | base::chisq.test | Yates' continuity correction | (Pearson 1900) |
| Anderson-Darling | kSamples::ad.test | Asymptotic p-value | (Scholz and Stephens 1987) |
| Kolmorgorov-Smirnov | base::ks.test | P-value approximate for discrete data, Two-sided p-value | (Kolmorgorov 1932) |
| *ClusterBurden* | base::fisher.test + base::chisq.test | Return 0 If either p-value equals 0 | (Fisher 1925) |
| CLUSTER | homepage.ntu.edu.tw/~linwy/CLUSTER.r | PARAMS=[num_perm = 1000, two-sided=TRUE] | (Cheung et al 2012) |
| DoEstRare | DoEstRare::DoEstRare | PARAMS=[alpha=0.05 , c=0.1] | (Persyn et al 2017) |
| WST | assoctesteR::WST | Asymptotic p-value | (Madsen and Browning 2009) |
| C-alpha | assoctesteR::CALPHA | Asymptotic p-value | (Neale et al 2011) |
| SKAT | assoctesteR::SKAT | Asymptotic p-value, PARAMS=[kernel="linear.weighted"] | (Wu et al 2011) |

*S3 Table: Type 1 error and power assessed via simulated data for proposed methods and published alternatives.*

| | *Clustering model:* | **Uniform** | | **One cluster** | | **Multiple clusters** | |
|---|---|---|---|---|---|---|---|
| | *Protein length:* | **500** | **1000** | **500** | **1000** | **500** | **1000** |
| **Cases** | **Unique variants** | 13.1 (5.8) | 26.1 (11.6) | 9.7 (6.2) | 19.4 (12.6) | 12.4 (6.3) | 24.8 (12.7) |
| **Controls** | **Unique variants** | 81.9 (82.2) | 163.9 (164.5) | 85.7 (85.9) | 171.1 (172.2) | 86.0 (86.3) | 172.3 (172.9) |
| | **Burden odds-ratio** | 2.49 (1.02) | 2.49 (1.03) | 1.71 (1.02) | 1.71 (1.03) | 2.28 (1.01) | 2.26 (1.02) |
| **Burden** | **Fisher-exact test** | 0.8 (0.042) | 0.97 (0.05) | 0.39 (0.044) | 0.58 (0.05) | 0.69 (0.042) | 0.87 (0.053) |
| **Two-sample goodness-of-fit tests** | ***BIN-test*** | 0.058 (0.051) | 0.058 (0.047) | 0.31 (0.051) | 0.52 (0.05) | 0.45 (0.05) | 0.73 (0.047) |
| | **Kolmorgorov-Smirnov** | 0.036 (0.023) | 0.039 (0.03) | 0.16 (0.024) | 0.31 (0.032) | 0.26 (0.026) | 0.49 (0.033) |
| | **Anderson-Darling** | 0.056 (0.051) | 0.054 (0.049) | 0.18 (0.05) | 0.31 (0.05) | 0.27 (0.05) | 0.48 (0.049) |
| **Position-informed RVATs** | ***ClusterBurden*** | 0.72 (0.05) | 0.94 (0.05) | 0.48 (0.049) | 0.7 (0.053) | 0.76 (0.047) | 0.93 (0.052) |
| | **DoEstRare** | 0.8 (0.058) | 0.96 (0.062) | 0.46 (0.06) | 0.64 (0.06) | 0.74 (0.058) | 0.9 (0.063) |
| | **CLUSTER** | 0.82 (0.054) | 0.97 (0.056) | 0.42 (0.055) | 0.61 (0.059) | 0.71 (0.053) | 0.88 (0.057) |
| **Standard RVATs** | **C-alpha** | 0.75 (0.051) | 0.94 (0.056) | 0.42 (0.053) | 0.59 (0.058) | 0.7 (0.051) | 0.87 (0.057) |
| | **WST** | *NA* (0.17) | *NA* (0.089) | *NA* (0.16) | *NA* (0.086) | *NA* (0.16) | *NA* (0.08) |
| | **SKAT** | *NA* (0.18) | *NA* (0.2) | *NA* (0.18) | *NA* (0.21) | *NA* (0.18) | *NA* (0.2) |

*S1 Methods: Origin and bioinformatics processing of the OMGL and HCMR datasets.*

A total of 5,393 cases of hypertrophic cardiomyopathy (HCM) were available by combing two datasets; the Oxford Medical Genetics Laboratory and the Hypertrophic Cardiomyopathy Registry (HCMR). There were 2,757 probands sequenced at OMGL, all referred by cardiac specialists between 2013 and 2018. The HCMR cohort consisted of 2,636 probands with a clinical diagnosis of HCM, also sequenced between 2013 and 2018.

Genetic analysis of OMGL cases consisted of; target enrichment with a custom-designed Agilent HaloPlex kit, sequencing on Illumina MiSeq, adaptor trimmed using Cutadapt and short or low quality read were discarded with Trimmomatic. The same genetic workflow was followed for the HCMR cohort with the exception of the target enrichment kit which was a custom-designed Illumina Truseq kit. Joint bioinformatic processing of both cohorts proceeded with mapping to the human reference genome (hs37d5 assembly) with BWA and haplotype calling and genotyping with an in-house pipeline adapted from the GATK Best Practices. All OMGL variants were confirmed by Sanger sequencing. All HCMR variants were visually confirmed by manual inspection of the BAM files.

*S2 Methods: Implementation details and properties of the forward-time simulator.*

### *Overview*

A forward-time simulation program was developed in the R programming language using object orientated programming with R's built-in reference classes. The data.table package (Dowle and Srinivasan 2019) was used to efficiently simulate populations. The simulator is capable of evolving protein sequences with differential selection across their linear form, imitating variant clustering observed in our hypertrophic cardiomyopathy genes. The simulator is sufficient to imitate real world datasets for the purpose of comparing power and type 1 error for different methods.

The demographic model, followed Kryukov *et al.* (2009) maximum likelihood model, and includes an emulation of recent European population history. The three-stage model begins with a burn-in stage of 10,000 generations with an initial diploid population of 8,100 individuals. A brief "bottleneck", acknowledging the out of Africa hypothesis, followed for 100 generations with a reduced population size of 7,900. Most importantly, there was a final stage of exponential growth allowing the population to reach a size of 900,000 individuals in 370 generations. Each generation evolved via a three-stage process of mutation, selection and mating.

### *Protein length*

To explore the relationship between protein length (number of amino acid residues) and statistical power, protein sequences were simulated at two different lengths; 500 and 1,000 residues. The chosen values for protein length fall within the range of observed protein lengths for both SWISSPROT (n=20,233, mean=560) and the HCM gene panel (n=35, mean=688; **Fig. 1**).

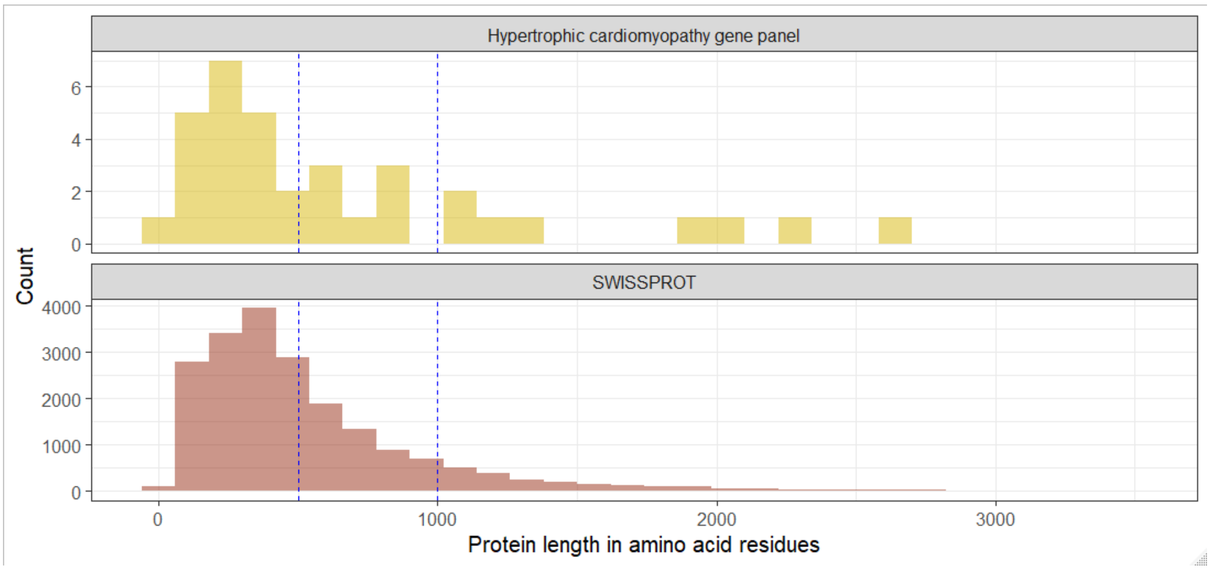

***Figure 1: Protein lengths in 20,233 SWISSPROT proteins and 35 genes associated with hypertrophic cardiomyopathy.*** *The x-axis is truncated at 3,500 residues, excluding very large proteins in SWISSPROT, such as TTN, that otherwise obscure the distribution. Blue dashed vertical lines at 500 and 1,000 denote the selected protein lengths for our simulations.*

*Mutations*

In each generation the number of new mutations was randomly drawn from a binomial distribution X~B(n,p) where *n* is the total number of mutable sites in the population and *p* is the mutation rate. The mutation rate was calculated by multiplying 1.8 x $10^{-8}$, the per-nucleotide base mutation rate in the human genome (Peng and Liu 2011), by the 3 bases in a codon and the empirical probability of a missense mutation 0.619. Thus, we made the simplifying assumption that all base changes are equally likely and all amino acids are present in equal proportions. This approximates an average missense mutation rate of 3.34 x $10^{-8}$.

To determine whether the simulated data was broadly similar to our observed data we compared the total number of segregating sites (SS) and the site frequency spectrum (SFS) to our HCM-gnomAD case-control dataset (**Fig. 2**). When SS were adjusted for sequence length and sample size, the number of SS were similar to the real data but had fewer on average. The SFS were comparable between the real and simulated data.

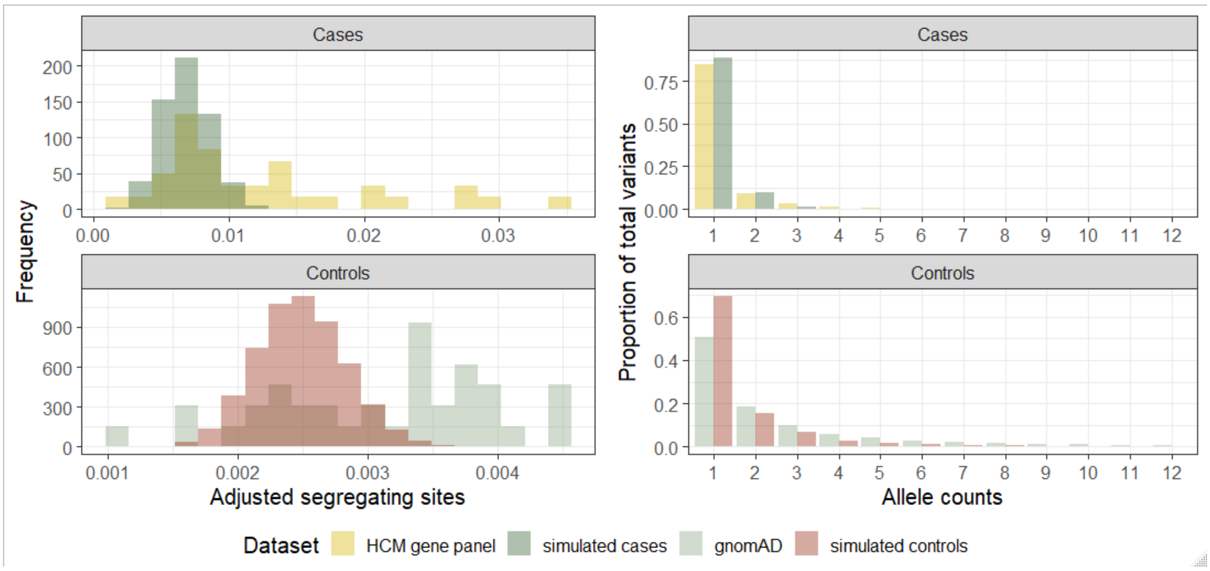

***Figure 2: The number of segregating sites and the site-frequency spectrum for simulated data and observed data.*** Segregating sites are adjusted for sample size and total number of sites in the sequence.

*Clustering models*

Three different amino-acid residue clustering models were explored; uniform, single-cluster and multiple-cluster. In the uniform model all variants in a protein are equally likely to be pathogenic. In the clustered models, only specific regions of the protein harbour pathogenic variants. Both clustering scenarios were apparent in the HCM gene panel data.

To implement the clustering scenarios, discrete pathogenic regions were specified before evolution. Cluster sizes were randomly drawn to span one-twentieth to one-quarter of the total protein length, were randomly located and could overlap. In the multiple-cluster model there was 70:30% chance of two or three pathogenic regions respectively. Each residue in the sequence was assigned a static selection coefficient (SSC). For the uniform model the SSC was 1 for every residue. For the clustered models, residues that fell within a pathogenic region (PR) were given SSCs of 5 and residues outside

PRs were given SSCs of 1. Normalized SSCs were calculated by dividing by the mean SSC across the protein so the resultant mean SSC of the sequence was 1. After evolution begins each new mutation was assigned a random selection coefficient (RCC) drawn from an exponential distribution (**Fig. 3**). Selection coefficients varied between 0.0001 and 0.1 where the log transformation of the distribution is uniform in this range. The SSC and the RCC were multiplied to give the final selection coefficient (FSC).

A non-random mating process was implemented whereby the chance of success was additively dependent on the FSC of each variant carried by an individual. As most variants were detrimental to survival, most carriers had reduced fitness and a reduced probability to mate. As the primary interest was in rare variants, recombination was ignored.

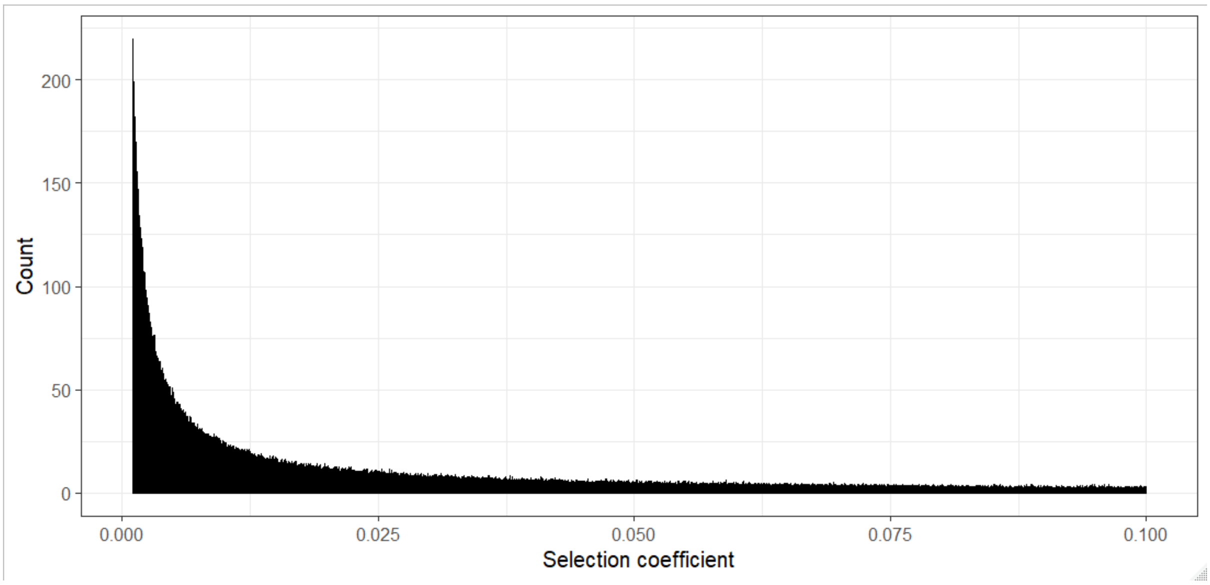

***Figure 3: The distribution of randomly allocated selection coefficients assigned to new mutations in the simulated populations.*** *The minimum coefficient was 0.0001 and the largest coefficient was 0.1.*

To monitor the selection procedure, FSCs were compared to the final allele counts of simulated mutations. There is an inverse relationship between allele count and mean selection FSC where high frequency variants had weaker FSCs (**Fig. 5; upper panel**). Variants within pathogenic regions, defined by the clustered models, had higher FSCs than variants outside these regions (**Fig. 5; lower panel**).

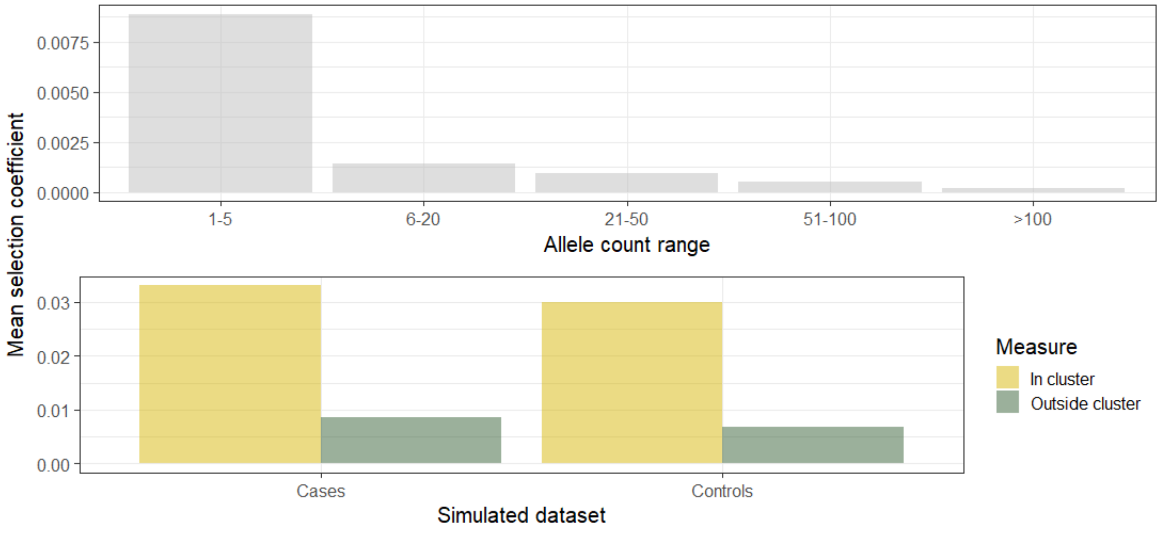

***Figure 5: Selection coefficients of simulated variants.*** *The upper panel shows the relationship between the final allele count of simulated variants and their static selection coefficients. The lower panel shows the average selection coefficient of variants that fall inside and outside the causal cluster(s).*

*Simulated datasets*

In order to generate simulated cohorts with differential distributions of variants, simulated phenotype was based on both the FSC and variant position. This is necessary as selection coefficient alone is insufficient to define a variant as disease-causing. This is analogous to an incompletely penetrant variant or a disease that does not lower reproductive fitness. Odds-ratios were drawn from an L-shaped distribution with mean 2.60 and median 2.35. All variants in the population were then ranked by FSC and assigned the equivalent ranked odds-ratio, therefore variants with the highest FSC had the highest odds-ratios. These odds-ratios were then multiplied by 3 or 1/3 depending on whether they fall within or outside a pathogenic region. Affection status was then simulated based on these odds-ratios according to the formula;

$$P(Affected|X) = \frac{e^{\alpha+BX}}{1+\ e^{\alpha+BX}}$$

Here $X$ is the vector of genotypes, $B$ is their respective odds ratios and alpha was defined by log(prevalence / 1 –prevalence). Prevalence was specified as 1/500 to correspond to the population prevalence of hypertrophic cardiomyopathy (Maron et al 1995). After each sample was assigned a phenotype, 2,500 cases and 123,000 controls were randomly selected from the population. For each gene length we simulated 10,000 populations in order to produce precise estimates of type 1 error with a binomial exact 95% confidence interval (500 successes in 10,000 trials) of 0.0458 to 0.0545.

*References*

Matt Dowle and Arun Srinivasan (2019). data.table: Extension of `data.frame`. R package version 1.12.2. https://CRAN.R-project.org/package=data.table.

Kryukov G., Shpunt A., Stamatoyannopoulos J., Sunyaev S. (2009). Power of deep, all-exon resequencing for discovery of human trait genes. PNAS. 106 (10): 3871-3876.

Peng B., Liu X. (2011). Simulating Sequences of the Human Genome with Rare Variants. Human Hereditary. 70; 287-291.

Maron B., Gardin J., Flack J., Gidding S., Kurosaki T., Bild D. (1995). Prevalence of hypertrophic cardiomyopathy in a general population of young adults. Echocardiographic analysis of 4111 subjects in the CARDIA Study. Coronary Artery Risk Development in (Young) Adults. Circulation. 92 (4): 785-9.

**S1 Figure: Distribution and risk predictions for rare-missense *MYH7* variants in our case-control cohort.**

The variant positions and training data for the GAM are a case cohort of 5,338 hypertrophic cardiomyopathy cases and 125,748 gnomAD controls. The density plot (lower panel) may give the impression that there is an excess of control variants in the C-terminus of the *MYH7* protein; however the GAM model (upper panel) resolves this potential misinterpretation and clearly shows an odds-ratio greater than 1 for the entire protein.

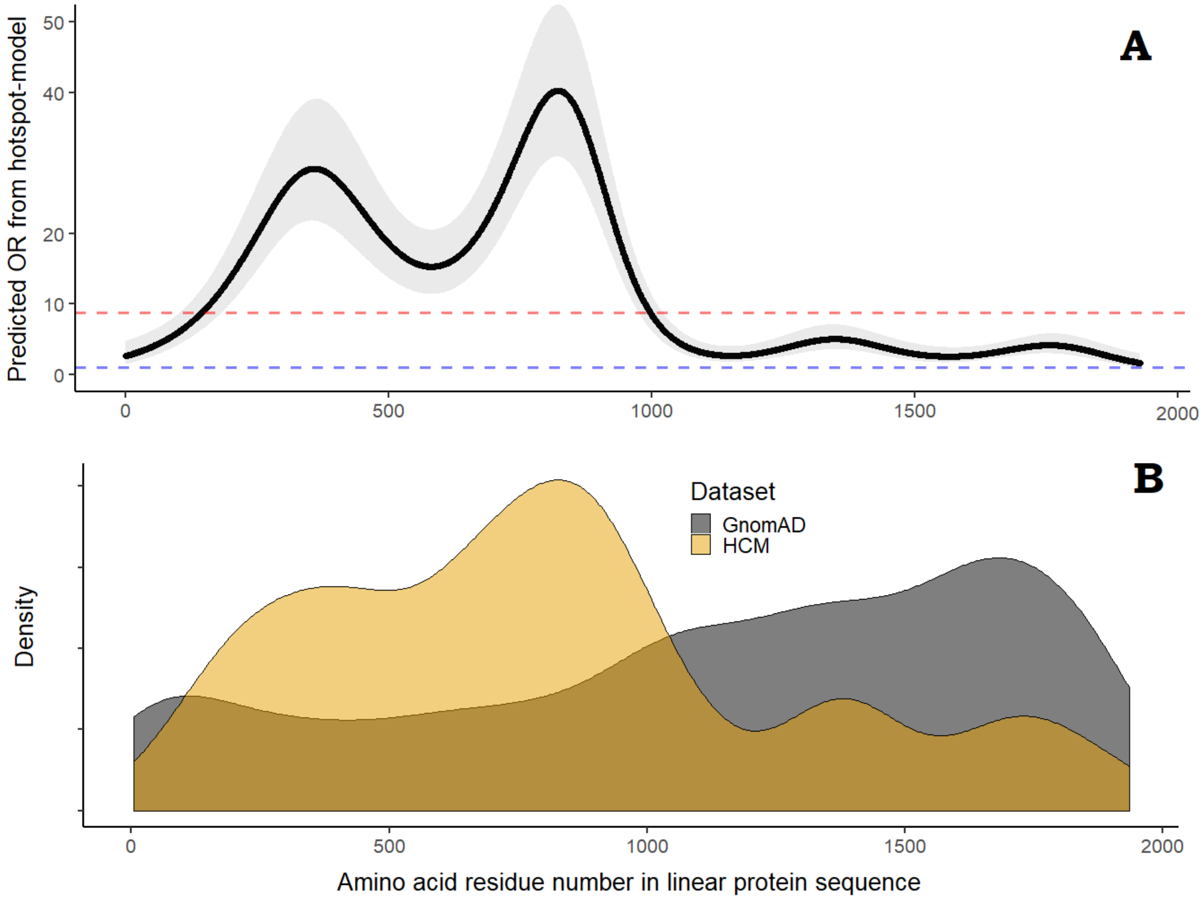

**S14 Figure: Rare-missense variant clustering in *MYBPC3* with and without potential founder mutations.**

Variant clustering model (*hotspot-model*) are generated for three different frequency filtering strategies. The model identifies four discrete regions with high pathogenic potential regardless of whether the founder mutations are included in the analysis. However, the magnitude of the predicted ORs are higher under normal filtering conditions (e.g. *popmax* < 0.01%).

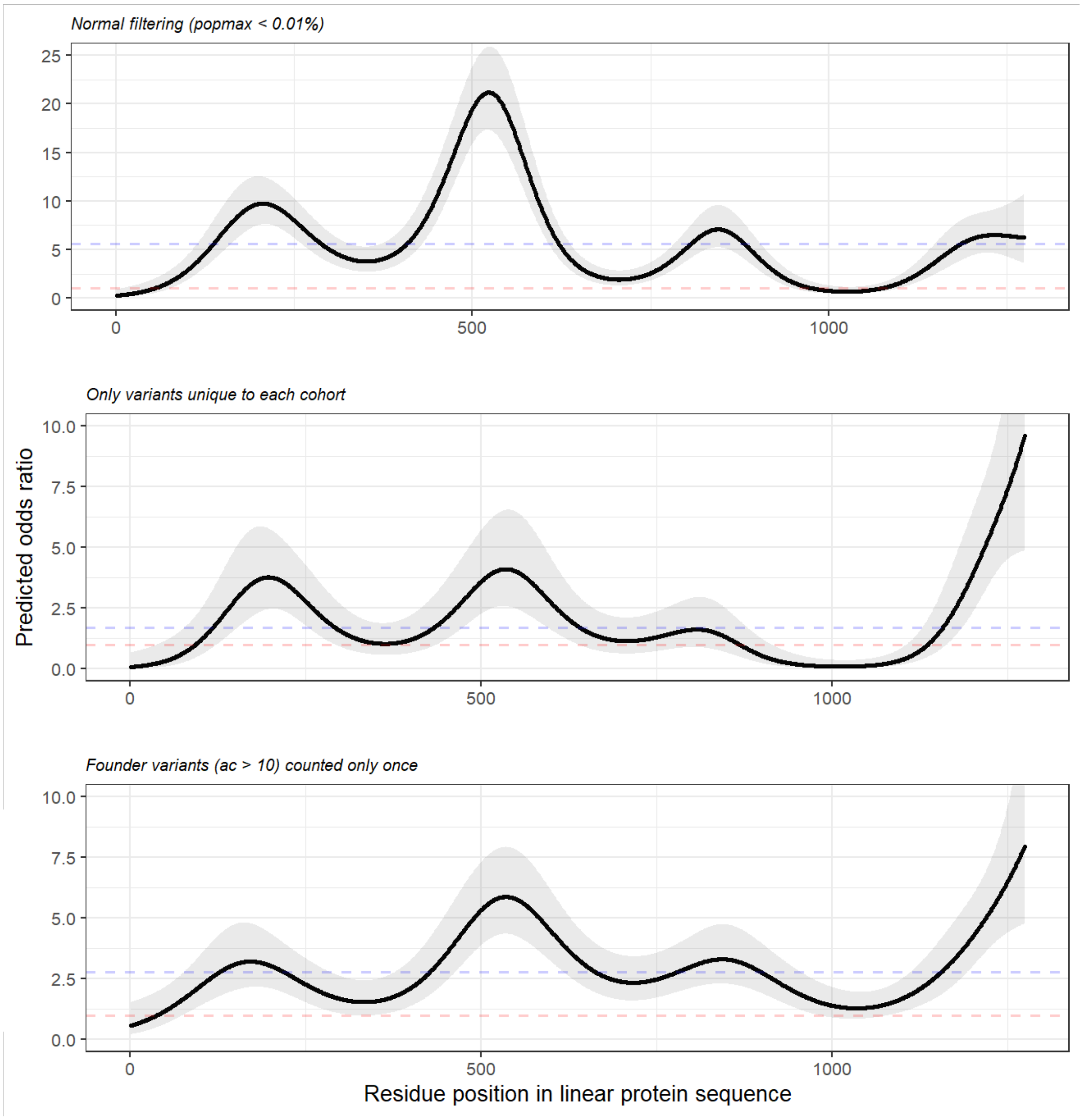
